# A biorefinery concept for the production of fuel ethanol, probiotic yeast and whey protein from a by-product of the cheese industry

**DOI:** 10.1101/2020.04.24.060889

**Authors:** María Dolores Pendón, José V. Madeira, David E. Romanin, Martín Rumbo, Andreas K. Gombert, Graciela L. Garrote

**Affiliations:** Centro de Investigación y Desarrollo en Criotecnología de Alimentos, CIDCA (UNLP-CONICET-CIC.PBA) La Plata. Argentina; University of Campinas, School of Food Engineering, Rua Monteiro Lobato 80, 13083-862, Campinas, SP, Brazil; Instituto de Estudios Inmunológicos y Fisiopatológicos (IIFP, UNLP-CONICET), La Plata, Argentina

**Keywords:** *Kluyveromyces*, whey, bioethanol, probiotics, biorefinery

## Abstract

1.

Agroindustrial by-products and residues can be transformed into valuable compounds in biorefineries. Here we present a new concept: production of fuel ethanol, whey protein and probiotic yeast from cheese whey. An initial screening under industrially relevant conditions, involving thirty *Kluyveromyces marxianus* strains, was carried out using spot assays to evaluate their capacity to grow on cheese whey or on whey permeate (100 g lactose/L), under aerobic or anaerobic conditions, in the absence or presence of 5% ethanol, at pH 5.8 or pH 2.5. The four best growing *K. marxianus* strains were selected and further evaluated in a miniaturized industrial fermentation process using reconstituted whey permeate (100 g lactose/L) with cell recycling (involving sulfuric acid treatment). After five consecutive fermentation cycles, the ethanol yield on sugar reached 90% of the theoretical maximum in the best cases, with 90% cell viability. Cells harvested at this point displayed probiotic properties such as capacity to survive the passage through the gastrointestinal tract and capacity to modulate innate immune response of intestinal epithelium, both *in vitro*. Furthermore, the CIDCA 9121 strain was able to protect against histopathological damage in an animal model of acute colitis. Our findings demonstrate that *K. marxianus* CIDCA 9121 is capable of efficiently fermenting the lactose present in whey permeate to ethanol and that the remaining yeast biomass has probiotic properties, enabling an integrated process for the obtainment of whey protein, fuel ethanol and probiotics from cheese whey.

**Importance:** Cheese whey is the liquid remaining following the precipitation and removal of milk casein during cheese-making. This by-product represents about 85-95% of the milk volume and retains 55% of milk nutrients so it can be exploited as a source of valuable end products. However, at a global level around 50% of cheese whey is wasted, representing an important environmental impact and indicating the need to develop alternative processes to recover value. *Kluyveromyces marxianus* is capable of fermenting lactose, generally regarded as safe, and has been explored separately as an ethanol producer and as a viable bioactive microorganism. The significance of our research is to establish the proof of concept that a biorefinery for fuel ethanol production using whey and *K. marxianus* can also be exploited to obtain viable probiotic biomass, conferring an added value to the process and providing an alternative to reduce environmental impact.

## 3. Introduction

Whey production around the world is over 160 million tons per year and it is continuously increasing (1). Whey is characterized by high ranges of biochemical oxygen demand (BOD) and chemical oxygen demand (COD), 30–50 g/L and 60–80 g/L, respectively, mainly due to its lactose content. In recent years, there has been a significant increase in alternative processes for the use of whey, which has contributed to reducing the environmental impact caused by the disposal of this by-product from the cheese industry. For every kilogram of cheese produced, roughly nine liters of whey (rich in lactose and protein, as well as some mineral salts) are generated on average, making this by-product a source of low-cost nutrients. Currently, large industries obtain this whey from the cheese industry and use it for the production of whey protein concentrate (WPC), which contains 35-80% protein, or with further purification processes it is possible to obtain whey protein isolate (WPI) with more than 90% protein, generating always as a leftover the so-called whey permeate, from which lactose can be recovered (2). However, depending on the country, region and production scale, there is still a high proportion of whey that is discarded with no treatment at all and even during the production of protein concentrate, a large amount of lactose-rich whey permeate is generated. In all these cases, ethanol production by lactose fermentation with appropriate yeast strains is an alternative to increase the value of the industrial waste and minimize environmental impacts (1).

Whey itself can be used as a substrate for fuel ethanol production (3); however, whey permeate also appears as a promising raw material, due to its high C/N ratio, which favors fermentative metabolism over cell growth (4) Furthermore, the use of whey permeate has the advantage of allowing the recovery of whey protein as a separate product, increasing the economic value of the whole process.

One important feature that yeast used for ethanol production from whey or whey permeate should have is the capacity to ferment lactose. Among these, *Kluyveromyces lactis*, *Kluyveromyces marxianus* and *Candida pseudotropicalis* are among the most studied species (5). *K. marxianus* shows interesting additional features for ethanol production from whey, such as thermotolerance, which allows the process to be carried out at higher temperatures, which in turn leads to decreased costs related to cooling, e.g. installation and operation of heat exchangers and use of cold water. Another interesting property is that this yeast presents a high specific growth rate, which confers a competitive advantage over contaminants in a non-aseptic industrial process. *K. marxianus* strains are generally recognized as safe (GRAS), mainly those isolated from food sources, such as kefir grains, a starter used to produce a fermented milk product (6, 7). Strains from this species present high resistance to acidity, antioxidant activity, tolerance to the gastrointestinal conditions and they also have the capacity to modulate inflammation of the gastrointestinal tract (7–9). Recovery of yeast biomass with probiotic features from fuel ethanol production using whey as a raw material could increase the added value of the whole whey recovery chain. Since the probiotic capacity of a microorganism may depend on physiological features and is conditioned, among other variables, by the substrate used for growing it (10), our aim was to establish a proof of concept that yeast with probiotic capacity can be recovered from a biorefinery for fuel ethanol production using whey permeate as the substrate. To this aim, we first screened several *K marxianus* strains from different origins for their capacity to grow on this substrate under anaerobic conditions, in the presence of ethanol and also at low pH, conditions that are relevant in an industrial process involving large fermentation volumes and cell recycling with sulfuric acid treatment to fight contaminants (11). Subsequently, we selected the best performing strains for evaluation in a miniaturized industrial process, after which viable biomass was recovered and tested for its probiotic properties both in vitro and in vivo.

## 4. Results

### 4.1. K. marxianus strains CIDCA 9121 and NCYC 1429 ferment whey permeate with 90% ethanol yield and high cell viability after 5 cycles in a scaled-down fermentation process

We screened the growth performance of thirty *K. marxianus* strains using whey (solidified with agar) as a substrate, in serial dilution spotting assays performed at 37 °C in a regular temperature cabinet (identified here as aerobic), for up to 24, 48 and 72 h. The Petri dishes contained whey reconstituted to reach 100 g/L of lactose. Depending on the experiment, 5% v/v of ethanol was added to the medium, pH was adjusted to 2.5 or plates were incubated under anaerobic conditions (Figure 1A). *K. lactis* CBS 2359, which is a well-known lactose-consuming species unable to grow in the absence of oxygen (12), and *S. cerevisiae* PE-2, which cannot grow on lactose, were used as controls. The strains were subsequently ranked according to the number of spots visible under each condition (Figure 1B) and ten of them, namely CIDCA 81111, CIDCA 81105, CIDCA 9121, NCYC 100, NCYC 2887, NCYC 2907, NCYC 3344, NCYC 3396, NCYC 1425 and NCYC 1429, were selected to be spotted on whey permeate-agar (Figure 2A). These strains were also ranked according to their growth capacity under the different conditions tested (Figure 2B). As a result, strains CIDCA 81111, CIDCA 9121, NCYC 2907 and NCYC 1429 were further selected for subsequent experimentation, due to their growth performance on whey permeate agar, especially under anaerobic conditions (Figure 2).

**Figure 1.**
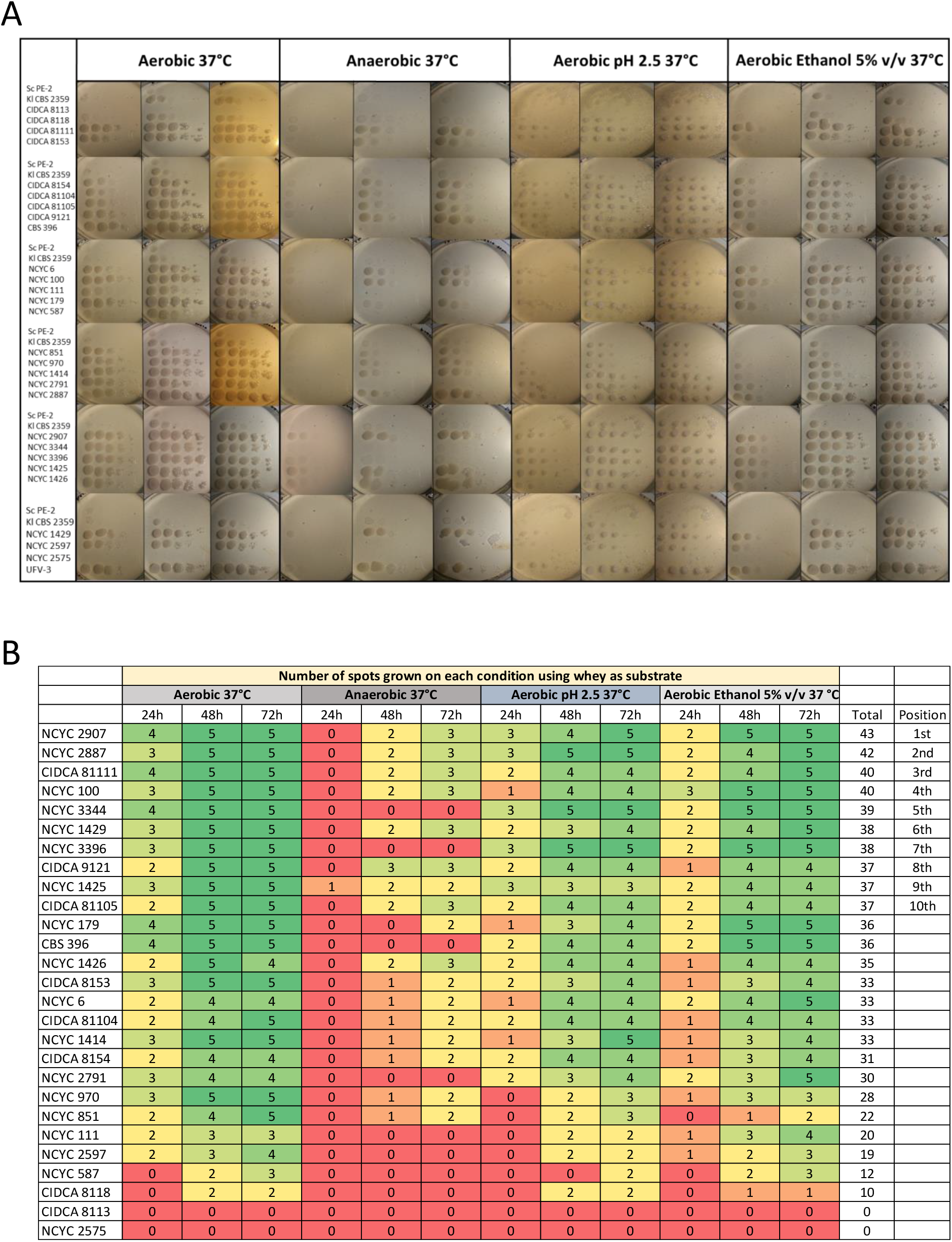
Spot tests of *K. marxianus* strains using whey as substrate. **A**: Growth of serial dilutions of different strains (from left to right, OD_600_: 10^−1^, 10^−2^, 10^−3^, 10^−4^ and 10^−5^) on whey-agar. The rows represent the different strains indicated in the figure and each column represents a specific condition: aerobic and anaerobic incubation, pH 2.5, and 5% v/v ethanol. Photographic records were made after 24 h, 48 h and 72 h of incubation at 37 °C. **B**: Quantitative scores of yeast strains’ growth capacity in the different conditions tested. Scores were obtained as the total number of spots for which growth at each dilution was evident.

**Figure 2.**
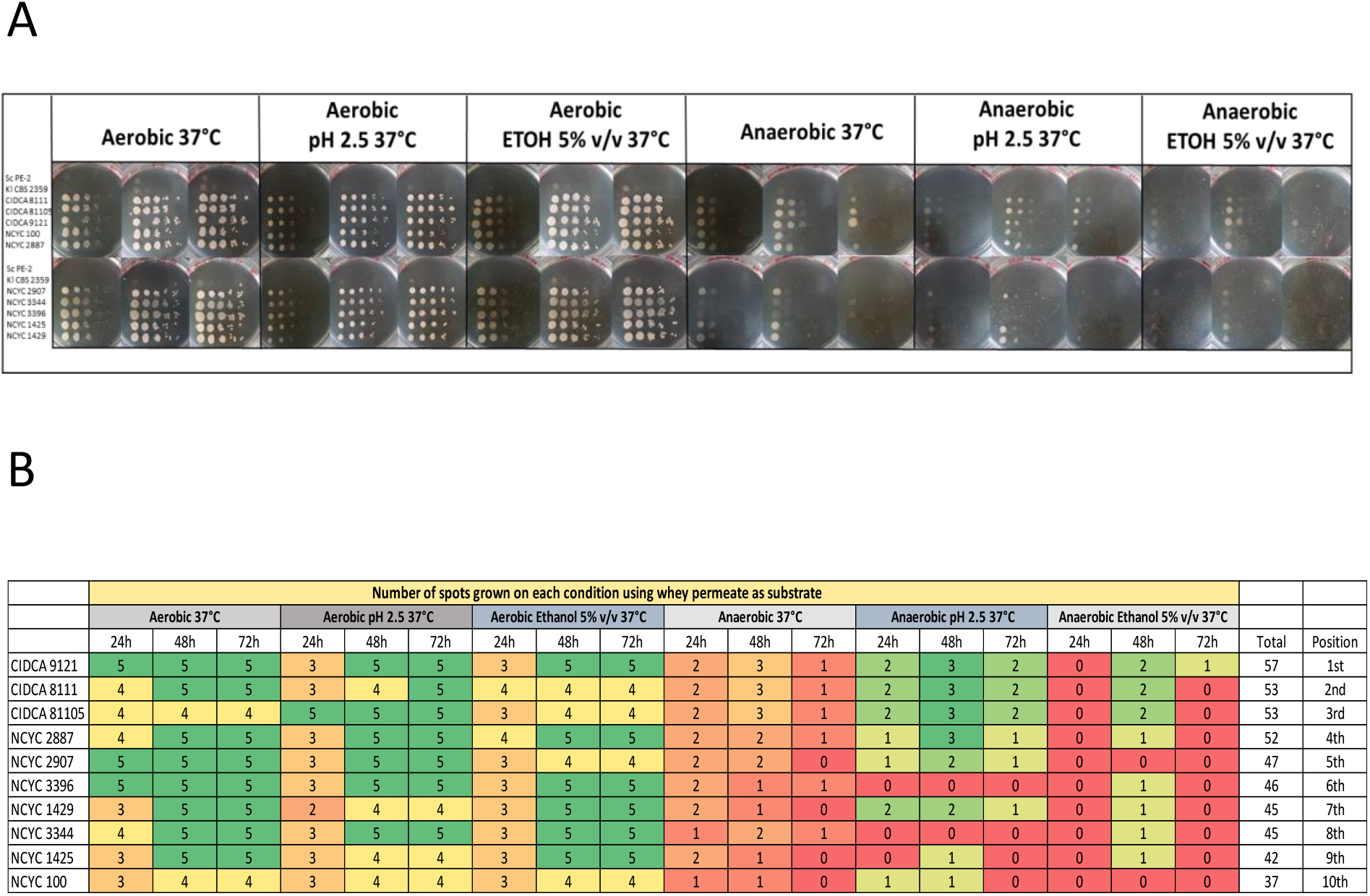
Spot tests of *K. marxianus* strains using permeate as substrate. **A**: Growth of serial dilutions of different strains (from left to right, OD_600_: 10^−1^, 10^−2^, 10^−3^, 10^−4^ and 10^−5^) in whey permeate agar. The rows represent the different strains indicated in the figure and each column represents a specific condition: aerobic and anaerobic incubation, pH 2.5, and 5% v/v ethanol. Photographic records were made after 24 h, 48 h and 72 h of incubation at 37 °C. **B**: Quantitative scores of yeast strains’ growth capacity in the different conditions tested. Scores were obtained as the total number of spots for which growth at each dilution was evident.

The next step consisted of cultivating the selected strains (individually) in a system previously validated as an adequate proxy for the industrial production of fuel ethanol from sugarcane molasses with cell recycling and sulfuric acid treatment, which is usually employed for fighting contaminating microorganisms in big scale non-aseptic fermentations(13). This system recapitulates the performance of a given yeast strain during consecutive fermentation cycles, a feature which cannot be captured using regular cultivation systems, such as flasks or tubes. The biomass formed at the end of one fermentation cycle is acid-treated and used as inoculum for the next fermentation cycle, and the key parameters to be followed along the cycles are cell viability and ethanol yield on sugars. All assayed strains showed the capacity to perform five consecutive cycles of fermentation in this system, with about 99% of lactose consumption in each 10 h cycle (Table 1). Depending on the strain considered, final ethanol concentration was around 3.5 to 4.0 % v/v. In Figure 3 (panels A and B), it is possible to observe that the ethanol yield varies between 80 and 91% of the theoretical maximum (0.511 grams ethanol formed per gram of hexose equivalent consumed), and that yeast viability remains high (above 89%) for two of the four investigated strains, *viz. K. marxianus* CIDCA 9121 and NCYC 1429. Besides, these two strains showed the lower cycle to cycle variations, and were selected for subsequent studies.

**Table 1.** Lactose utilization and ethanol production using recycled *Kluyveromyces marxianus* strains during successive fermentation cycles

| Strain | Initial lactose<br>(g.L<sup>-1</sup>) | Residual lactose<br>(g.L<sup>-1</sup>) | Ethanol produced<br>(g.L<sup>-1</sup>) * | Time<br>(hours) |
| --- | --- | --- | --- | --- |
| CIDCA 81111 | 111.6 | 1.38 ± 0.02 | 35.5 ± 0.5 | 10.0 |
| CIDCA 9121 | 111.6 | 1.28 ± 0.01 | 39.7 ± 0.5 | 10.0 |
| NCYC 2907 | 111.6 | 1.13 ± 0.01 | 34.8 ± 0.3 | 10.0 |
| NCYC 1429 | 111.6 | 1.09 ± 0.03 | 39.5 ± 0.5 | 10.0 |

The values denote the mean of a biologic duplicate ± standard error of the mean (± SEM)
\* Maximum ethanol concentration detected trough the fermentation cycles

**Figure 3.**
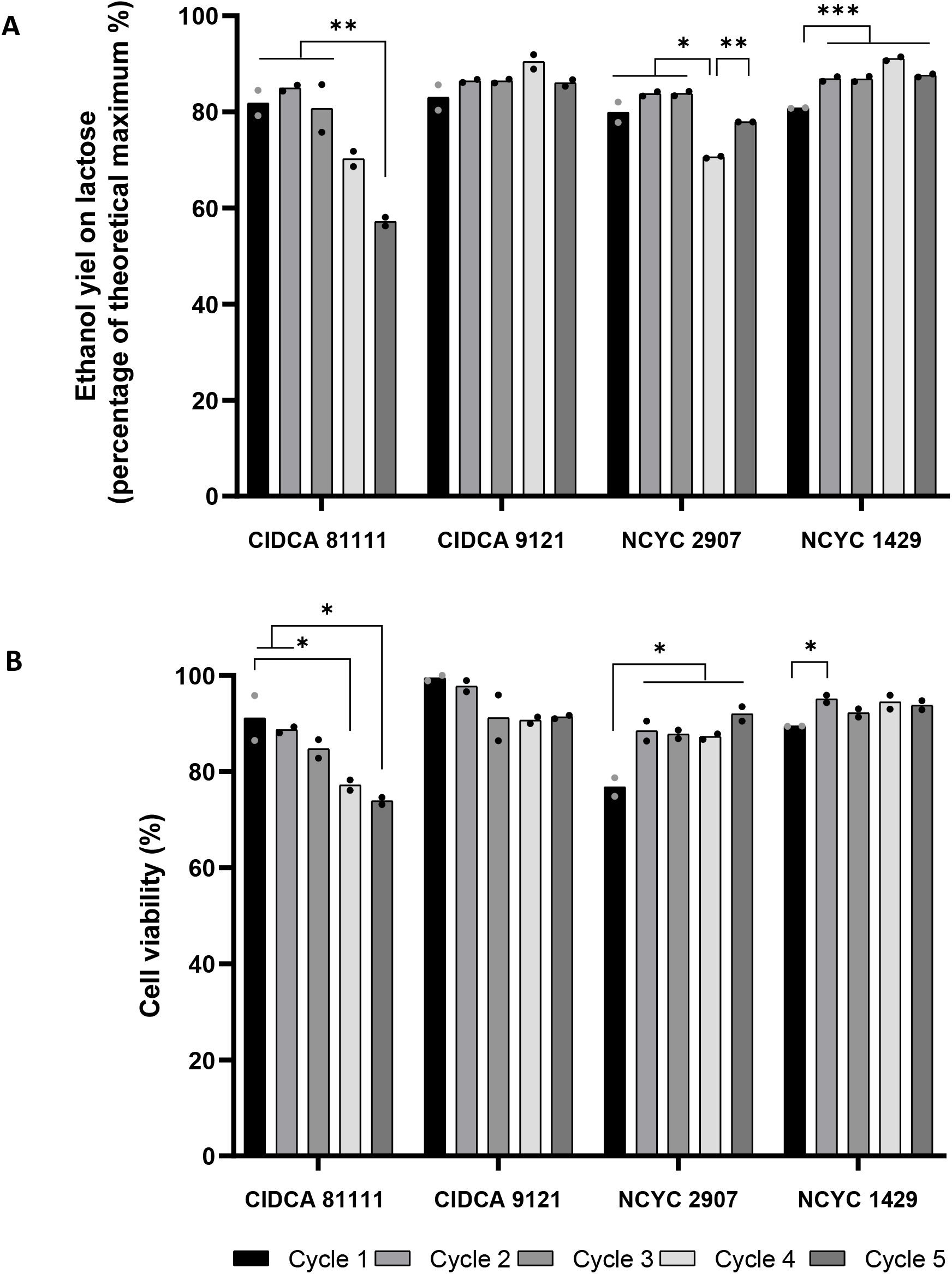
Ethanol yield (A) and cell viability (B) of *K. marxianus* CIDCA 81111, CIDCA 9121, NCYC 2907 and NCYC 1429 after each cycle of whey permeate fermentation at 34 °C in a miniaturized system. **A:** Ethanol yield as a percentage of the theoretical maximum (0.51 g ethanol.g hexose-equivalents^−1^) for the four strains during five fermentation cycles. **B:** Cell viability over the five cycles for the four yeast strains, measured as the percentage of viable cells in a population of viable and nonviable cells. Mean value and individual independent biological replicates are depicted. Cycle to cycle differences were evaluated using a One-way ANOVA test with Dunnett multiple comparison test * *p* < 0.0332; ** *p* < 0.0021; *** *p* < 0.0002.

To verify whether temperature (30 °C or 37 °C) and/or medium composition (whey or whey permeate) could have any influence on process performance, the ethanol yield and cell viability were measured after each of five consecutive fermentation cycles, for the two selected strains (Figure 4). For both strains, fermentation at 37 °C resulted in a ~10% higher ethanol yield on lactose, when compared to the performance at 30 °C (Figure 4, panels A and C *p* < 0.001). Likewise, at 37 °C the yield in ethanol was higher using permeate than serum as substrate also for both tested strains (Figure 4, panels A and C *p* < 0.001). Furthermore, cell viability was less influenced than ethanol yield by the temperature (Figure 4, panels B and D), showing only an increased viability in the case of strain CIDCA 9121 growing on whey at 37 °C (Figure 4, panel B *p* < 0.05). Consequently, whey permeate fermentation at 37°C was selected as the condition for subsequent experiments.

**Figure 4.**
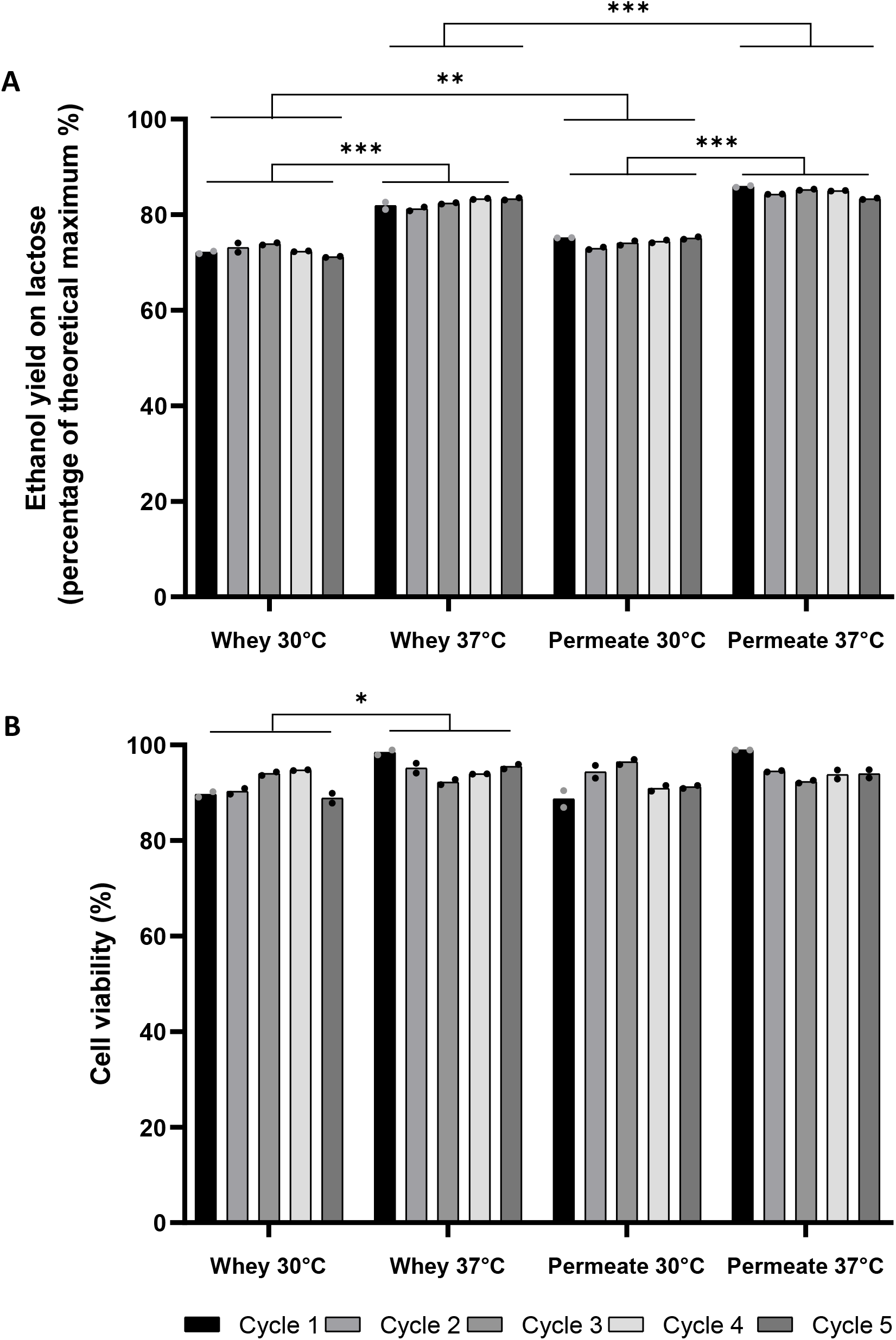
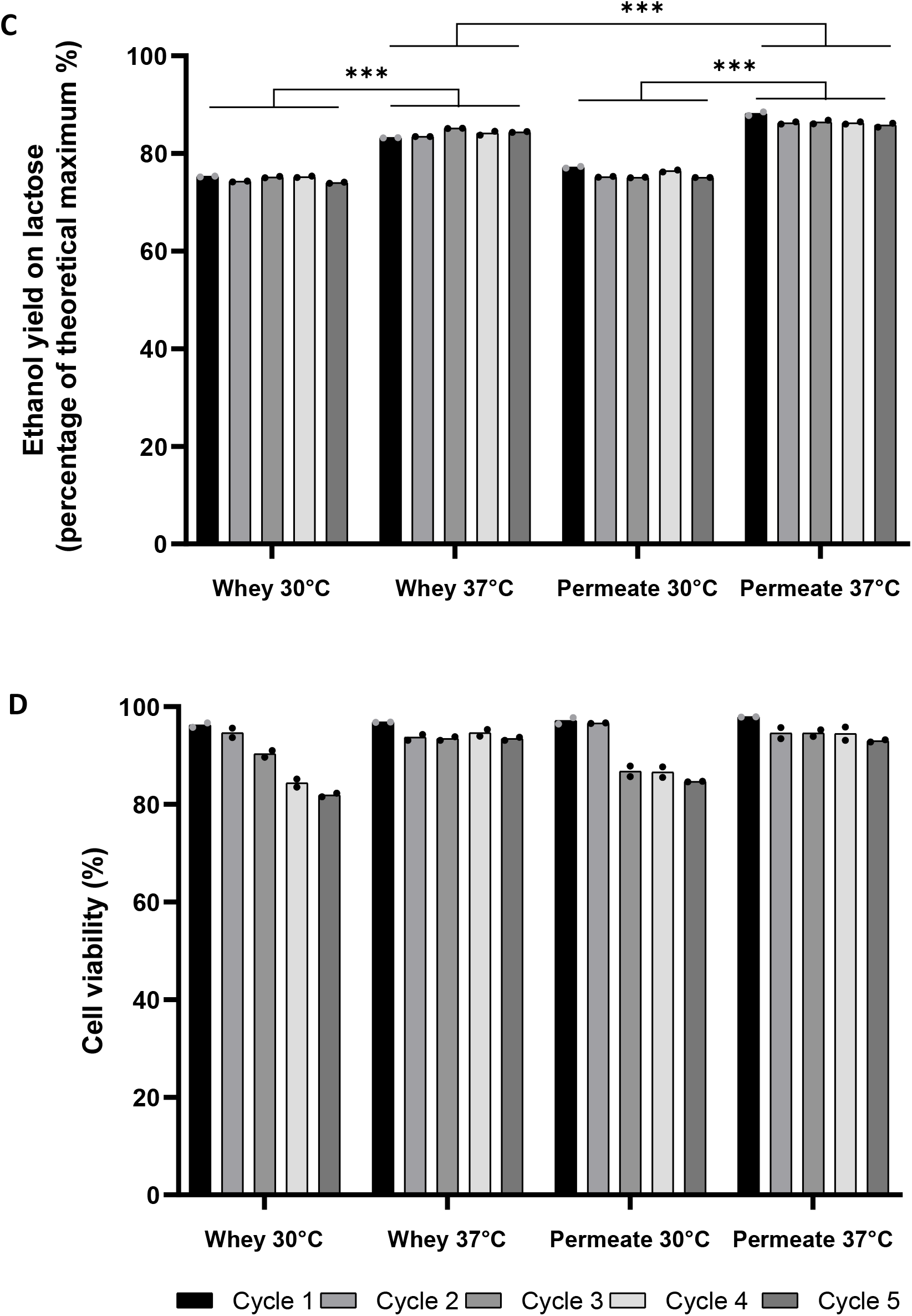
Ethanol yield and cell viability of *K. marxianus* CIDCA 9121 (A, B) and NCYC 1429 (C, D) after each cycle of fermentation at 30 and 37 °C using whey and permeate as substrate in a miniaturized system. **A, C:** Ethanol yield as a percentage of theoretical maximum (0.51 g ethanol.g hexose-equivalents^−1^) for the two strains during the five fermentation cycles. **B, D:** Cell viability over the five cycles for the two yeast strains, measured as the percentage of viable cells in a population of viable and nonviable cells. Mean value and individual independent biological replicates are depicted. Paired comparison of performance obtained under different conditions was evaluated by T-test with Mann Whitney Test * *p* < 0.0332; ** *p* < 0.0021; *** *p* < 0.0002.

Since the original protocol published by Raghavendran *et al* (2017) (13) involved fed-batch fermentations (using 3 installments), we also evaluated the effects of using simple (or batch) fermentation instead. As may be observed from the results of *K. marxianus* CIDCA 9121 in Figure 5, cell viability remained unaffected and a slight improvement of the overall ethanol yield (*p* < 0,001) was detected, indicating the process can also be carried out using the much simpler batch mode.

**Figure 5.**
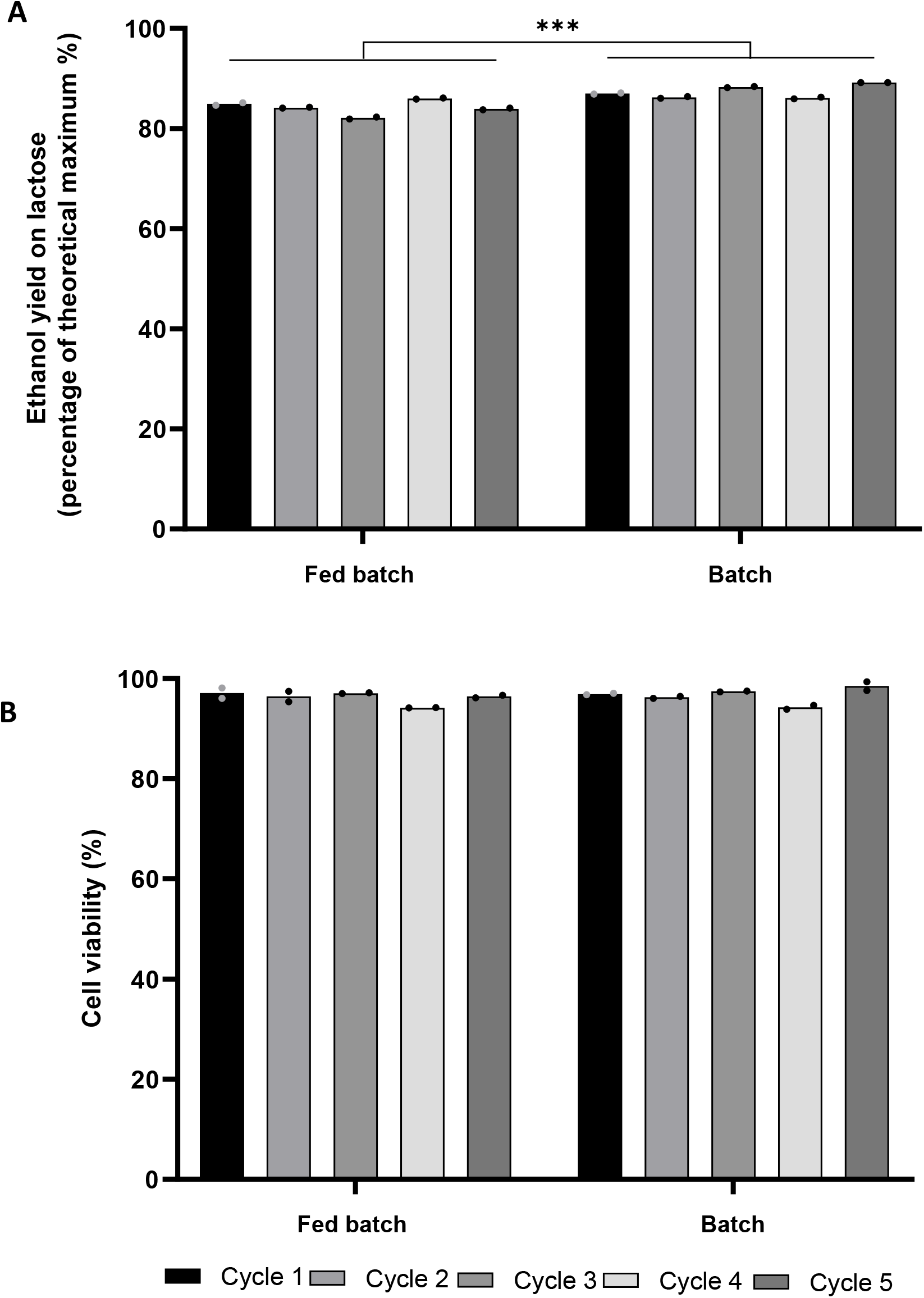
Ethanol yield (A) and cell viability (B) of *K. marxianus* CIDCA 9121 after each cycle of fermentation using permeate as substrate in simple batch and fed-batch processes. **A:** Ethanol yield as a percentage of theoretical maximum (0.51 g ethanol.g hexose-equivalents^−1^) during the five fermentation cycles. **B:** Cell viability over the five cycles measured as the percentage of viable cells in a population of viable and nonviable cells. Mean value and individual independent biological replicates are depicted. Paired comparison of performance obtained under different conditions was evaluated by T-test with Mann Whitney Test. * *p* < 0.0332; ** *p* < 0.0021; *** *p* < 0.0002.

### 4.2. Selected K. marxianus strains are capable to resist GIT conditions in vitro or in vivo and do not translocate to systemic compartment

To be used as a probiotic, one main feature that the selected strains should have is the capacity to survive the passage through the gastrointestinal tract. According to the results from the first screening on plates, described above, we decided to evaluate *K. marxianus* strains CIDCA 9121, NCYC 1429, CIDCA 81111 and NCYC 2907 for this capacity. All four strains showed good tolerance to gastrointestinal conditions *in vitro*, since the decrease of only one log (strains CIDCA 81111 and CIDCA 9121) or 1.5 log (strains NCYC 2907 and NCYC 1429) in the concentration of viable yeast cells was observed (Figure 6).

**Figure 6:**
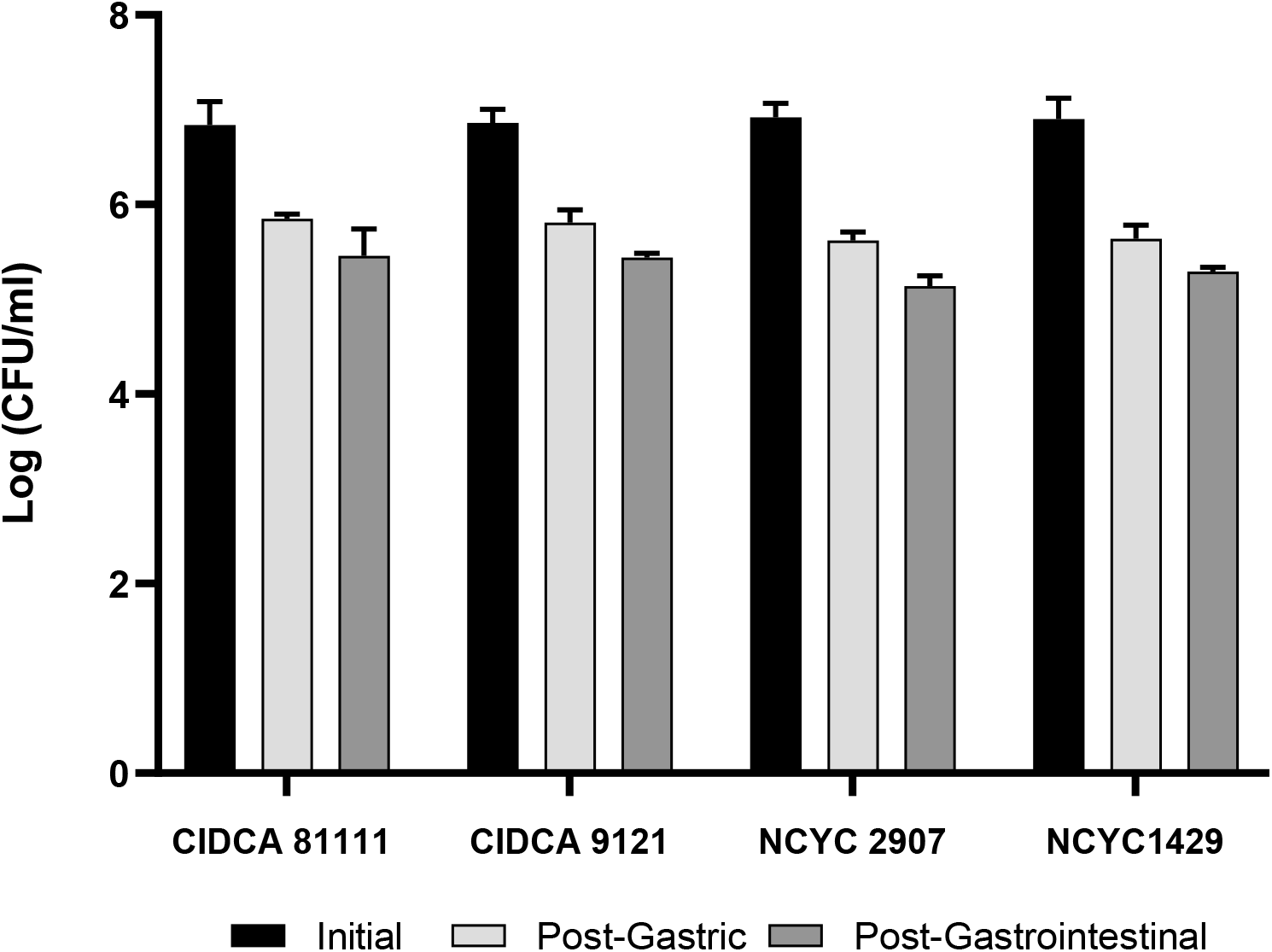
Viability of *K. marxianus* strains after five consecutive cycles of fermentation in a miniaturized industrial process, further subjected to a simulated gastrointestinal treatment. Survival of CIDCA 81111, CIDCA 9121, NCYC 2907 and NCYC 1429 after passage through simulated gastric (light grey) and subsequent intestinal (grey) conditions. Initial bacterial counts are shown in black bars. Results are expressed as mean ± SEM of two independent experiments. CFU/mL: colony forming units per milliliter. *P < 0.05 vs initial bacterial count; **P < 0.05 vs gastric conditions. Differences were evaluated using a One-way ANOVA test with Dunnett multiple comparison test * *p* < 0.05; ** *p* < 0.01; *** *p* < 0.001.

A suspension of 1.33 ± 0.95 10^7^ UFC/mL of CIDCA 9121 obtained after five consecutive cycles of fermentation simulating the industrial process was administered to mice. *In vivo* experiments confirmed their capacity to resist gastrointestinal conditions as they were recovered in mice feces (n = 5) at a concentration of 1.12 ± 0.44 10^7^ UFC/g, 2.22 ± 1.9 10^7^ UFC/g and 2.26 ± 0.8 10^7^ UFC/g after two, three and four days of oral administration, respectively. When yeast distribution in mice intestines was studied, a high concentration at the distal portions (ileum and colon) were observed for the two strains (not shown). Additionally, an important safety issue was that both strains did not show the capacity to translocate from the gastrointestinal tract to systemic compartments, since no yeast were recovered from liver or spleen.

### 4.3. Strain K. marxianus CIDCA 9121 cultivated in a miniaturized industrial fermentation protects against histopathological damage induced by TNBS in an acute animal model of colitis

One important aspect of probiotic yeast is their capacity to modulate the innate immune response. Since probiotic features may depend on the physiological status of the microorganisms, their growing on different substrates may affect their properties. Thus, we tested if the different yeast strains grown under industrially relevant conditions were able to modulate the innate immune response. The technique used to determine anti-inflammatory activity involves the use of genetically modified intestinal cells (Caco-2 ccl20: luc), containing the CCL20 promoter-controlled luciferase gene, which in turn responds to stimulation by the inflammatory agent FliC. Therefore, there is a direct correlation between increased luciferase activity and increased inflammatory response, as described by Nempont *et al* (2008) (14).

Four selected yeast strains were tested in this system. All of them displayed a strong capacity to reduce the Caco-2 cell activation induced by FliC to almost the basal levels (Figure 7A, *p*< 0.001). It is important to point out that the anti-inflammatory capacity was maintained after 2 and 4 fermentation cycles in the miniaturized industrial setting, which we tested with strain CIDCA 9121 (Figure 7 B).

**Figure 7.**
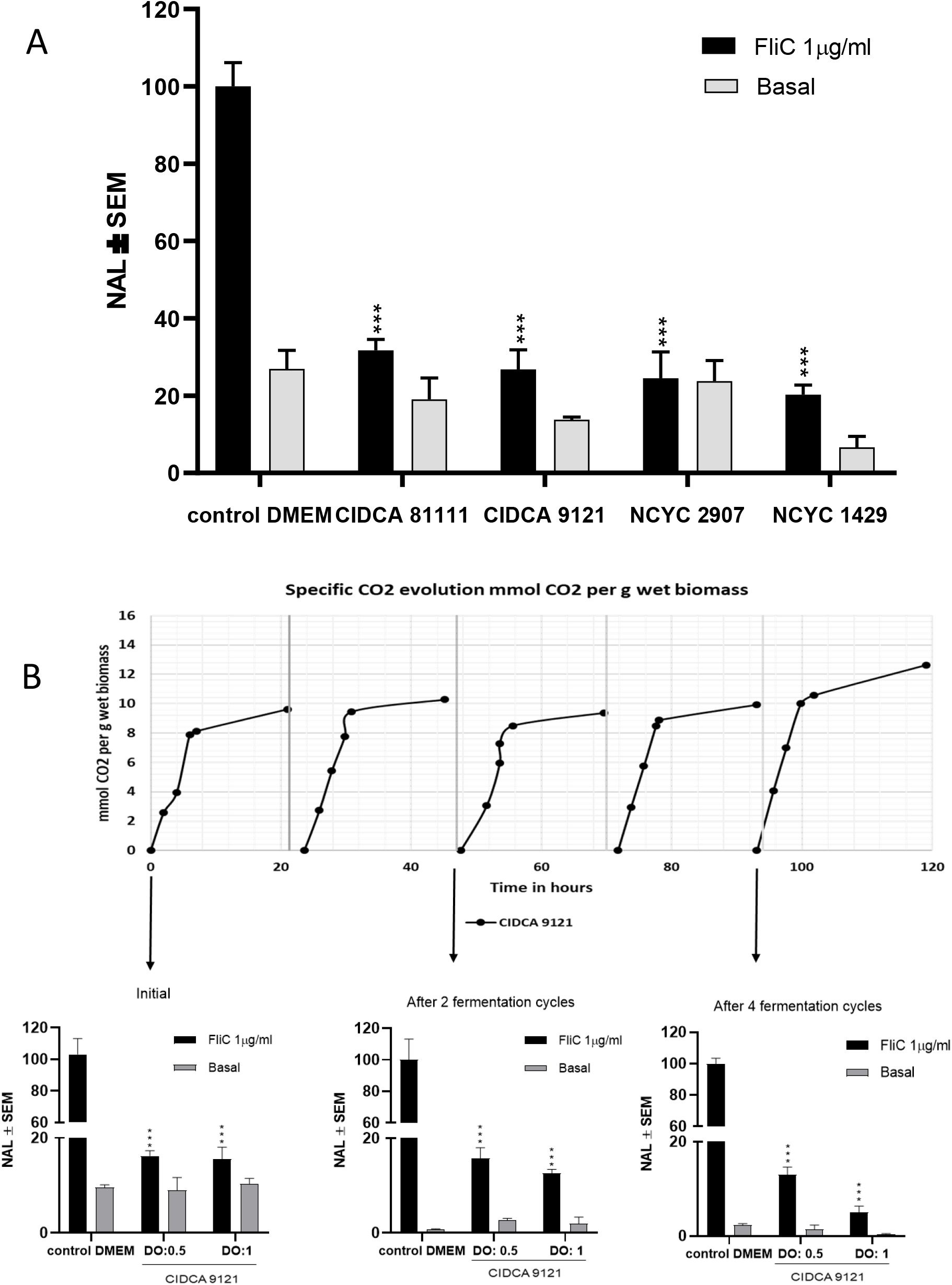
A: Pre-treatment with yeasts obtained after whey permeate fermentation leads to downregulation of inflammatory response in Caco-2-CCL20:luc cells. Reporter cells were stimulated with flagellin (1 μg/mL), after pretreatment with yeasts. Luciferase activity was determined in a cell lysate 6 h after stimulation. Results representative from three different experiments are shown. **B: Downregulation of inflammatory response in Caco-2-CCL20:luc cells by *K. marxianus* CIDCA 9121 after 0, 2 and 4 cycles of whey permeate fermentation at 37 °C in a miniaturized system**. Fermentation progression in each cycle was monitored by CO_2_ generation. Results are expressed as normalized luciferase activity (NAL), using the levels of stimulated cells in the absence of yeast cells as 100% of activation. Results shown are the mean from independent triplicates. Differences were evaluated using a One-way ANOVA test with Tukey multiple comparison test. *** Indicates a significant difference from the cells without treatment and stimulated with flagellin, with *p* < 0.0002.

To assess the ability of strain CIDCA 9121 obtained from the miniaturized industrial setting to mitigate tissue damage on an acute colitis model, BALB/c mice were administered a suspension of the yeast obtained after five cycles of fermentation on whey permeate in drinking water. Yeast suspension was prepared fresh on a daily basis. Control mice received normal water and both groups received normal food during the experience. After 24 and 48 h of TNBS instillation, animals showed a decrease in body weight (not shown). Yeast-treated animals showed more moderate weight loss. This difference is correlated with the histopathological damage in colon, which is the usual “gold standard” to evaluate protection in this model. Animals that received the yeast treatment showed less damage in epithelium and less infiltration in the submucosa compared to the TNBS/water group (*p* < 0.04, Figure 8). These results indicate that the administration of CIDCA 9121, even after being grown on simulated industrial ethanol production conditions, protects against histopathological damage induced by TNBS in an acute animal model of colitis (*p* < 0.05).

**Figure 8.**
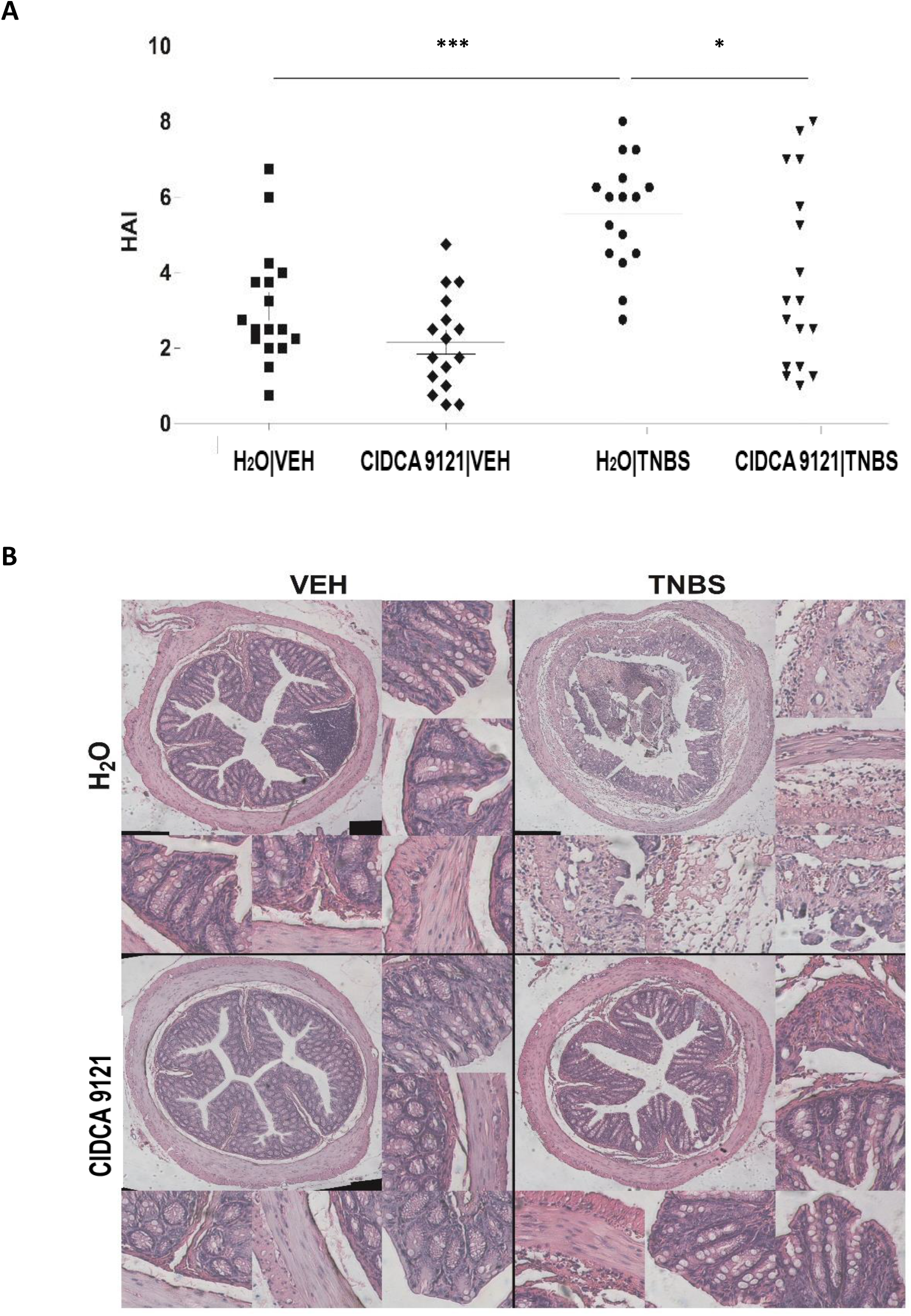
Oral administration of *K. marxianus* CIDCA 9121 protects animals against damage in TNBS acute colitis model. Mice were treated with yeast suspensions or water 4 days before TNBS or vehicle instillation and 48 h later tissue was collected for histopathological analysis. Results from n of 18 animals per group are shown. **A:** Histopathological activity index assigned to different experimental groups. Significant differences between groups are indicated in the graph. **B:** Photomicrograph of H&E-stained cross section of distal colon of a representative mouse of each experimental group. Paired comparison of performance obtained under different conditions was evaluated by T-test with Mann Whitney Test. * *p* < 0.0332; ** *p* < 0.0021; *** *p* < 0.0002.

## 5. Discussion

Cheese whey permeate is a by-product of the dairy industry that is partly used for lactose extraction, for the main purpose of being added to baby milk formulas. However, there is still a large volume of permeate worldwide that needs to be disposed of, representing cost and environmental issues. In this context, a process that could convert this permeate into fuel ethanol would be a valorization alternative. The use of permeate, instead of whey, does not affect ethanol yields, as demonstrated here (Figure 4), meaning that the protein found in whey can still be recovered before fermentation. On the other hand, the fermentation process naturally involves some yeast growth, meaning that in every cycle part of the cell biomass needs to be drained, in order to keep the microbial population below adequate thresholds. We hypothesized that this biomass could be used as a probiotic, rendering a third product in the biorefinery concept proposed here (15). However, although it had already been demonstrated that some *K. marxianus* strains present probiotic features (7, 9), it was essential to demonstrate that the yeast cells, after being used in the fermentation process to produce ethanol, would retain viability and probiotic activity.

Due to the high phenotypic polymorphism inherent to the *K. marxianus* species (16), this study started by screening 30 strains for their capacity to grow on plates containing whey (solidified with agar) under conditions relevant in the context of industrial fuel ethanol production, such as anaerobiosis, low pH and high ethanol concentrations (17). From this initial screening, we selected ten strains for similar experiments, but now using whey permeate, instead of whey. From this second screening round, four strains were selected for their evaluation in a miniaturized system that resembles the industrial production of fuel ethanol in Brazil from sugarcane, in terms of ethanol yield on lactose and cell viability. This system was originally developed and validated for the fermentation of diluted sugarcane molasses with the yeast *Saccharomyces cerevisiae*, with cell recycling and sulfuric acid treatment of the yeast biomass after each fermentation cycle, in order to reduce the contamination load, similarly to industrial practice (13). As a result, it was demonstrated that only after some fermentation cycles (typically five), the superior phenotype of some selected industrial strains becomes evident. We adapted this system to our study by using reconstituted whey permeate as a substrate and *K. marxianus* as the fermenting microorganism.

The concentration of lactose used here was ~100 g/L, in contrast to the typical ~200 g/L total reducing sugars concentration found in sugarcane-based media, mainly because this sets a limit to the concentration of ethanol along the fermentation (~50 g/L), which is important because of the lower tolerance displayed by *K. marxianus* to this alcohol, when compared to *S. cerevisiae* (11). This lower concentration of ethanol has negative consequences on the downstream distillation operations, but it has been claimed that this unit operation requires in general a minimum of 40 g/L ethanol to be economically viable (18).

Raghavendran *et al* (2017) (13) approximated the industrial fed-batch mode of operation by adding the medium in 3 installments along each fermentation round, which is mainly to avoid high sugar concentrations and the undesired effects of glucose repression that are known to affect *S. cerevisiae*. Since in our study lactose is the sugar present in the medium, meaning that no glucose should build up in the extracellular environment (lactose metabolism in *Kluyveromyces* yeasts takes place via its transport into the cells and subsequent intracellular hydrolysis into glucose and galactose (19), we decided to verify whether a simple batch process could be used in our experiments. As shown in Figure 5, batch operation leads to very similar results, when compared to fed-batch operation, meaning that the large-scale process could be simplified to batch mode, leading to lower costs and lower risk of failure. Another important aspect regarding industrial operation relates to the fact that fermentation performance at 37 ° C was better than at 30 ° C, in terms of both cell viabilities and ethanol yields. This means lower costs related to fermenter cooling, in terms of equipment purchase and maintenance, and cold-water usage.

The main aim of this work was to establish the proof of concept that yeast with probiotic features can be obtained after fermentation on a by-product from the dairy industry, namely the conversion of lactose into ethanol. Probiotics are viable microorganisms that when administered in the appropriate amount confer a health benefit to the consumer (20). The health benefit can be related to different properties such as the capacity to complement a metabolic activity of the consumer, the capacity to block the infectious cycle of a given pathogen or the modulation of local or systemic immunity that may confer the capacity to fight an infection or avoid excessive inflammation, among others. Different microorganisms have shown probiotic features, being remarkable that these properties in most of the cases are strain-dependent. Among yeasts, strains from the *Saccharomyces* and *Kluyveromyces* genus have been identified as probiotics (21, 22). Interestingly, *K marxianus* has shown capacity to reinforce the epithelial barrier and improve performance in animal production (23, 24). We have identified *K. marxianus* strains isolated from kefir grains with the capacity to also modulate inflammation and oxidative stress *in vivo* (9). Many of these features may depend on the expression of surface or cytoplasmic determinants responsible at the molecular level for the observed phenotype. It has been shown that probiotic lactic acid bacteria may change their probiotic capacity when grown on different substrates (10). Furthermore, extensive transcriptomic changes have been described for *K. marxianus* under 6% ethanol stress (25). Even if this was not the aim of the referenced work, it can be speculated that these transcriptomic changes could affect probiotic features of ethanol-stressed yeasts. Nevertheless, our results indicate that *K. marxianus* CIDCA 9121 conserves its probiotic features after bioethanol production, among them the capacity to modulate the inflammatory response in *in vitro* and *in vivo* models. These results establish the proof of concept that a biorefinery for ethanol production using whey or whey permeate can also be exploited to obtain viable probiotic biomass, conferring an added value to the process. The sustainability of biorefineries depends on the efficiency to use resources and minimize waste products which may rely on the integration of different processes (26–28). The process we studied in the present work is in line with this concept aiming at using a food industry waste as substrate to obtain fuel ethanol, protein concentrate and probiotic biomass.

We have shown the feasibility of keeping probiotic features upon growing on industrial waste under conditions similar to industrial fermentations, opening the possibility to generate probiotic biomass as a way to add value to the whole process (15). There are several possibilities of implementation of a biorefinery as proposed here, since whey or whey permeate can be concentrated for transport logistic purposes and also to achieve different final levels of ethanol during the fermentation. The different operations should be performed with a high threshold of microbiological security to avoid incorporation of microorganisms that may spoil the fermentation process. Under these conditions, it should be feasible to implement a “food grade” probiotic yeast production in a bioethanol biorefinery, adding a new option for the renewable energy industry.

## 6. Material and methods

### 6.1. Yeasts strains and maintenance

*Kluyveromyces marxianus* strains from different culture collections (Table 2) were first cultured in YPD medium (yeast extract 10 g/L, bacteriological peptone 20 g/L, and glucose 20 g/L) at 30 °C for 24 h in 20 ml tubes with 5 ml of medium without stirring. Frozen stock cultures were stored at −80 °C in YPD-medium with 50% (v/v) glycerol.

**Table 2.** Yeast strains used in this work

| Species | Number of strains | Identification * | Relevant properties | Reference |
| --- | --- | --- | --- | --- |
| *Kluyveromyces marxianus* | 18 | NCYC 6, NCYC 100 NCYC 111, NCYC 179 NCYC 587, NCYC 851, NCYC 970, NCYC 1414, NCYC 2791, NCYC 2887, NCYC 2907, NCYC 3344, NCYC 3396, NCYC 1425, NCYC 1426, NCYC 1429, NCYC 2597, NCYC 2575 | Ethanol production, anaerobic growth | Lane et al.(32) |
|  | 2 | CBS 396, CBS 712 |  | Signori et al.(33) |
|  | 1 | UFV-3 | Ethanol production, sequenced genome | Silveira et al.(34) |
|  | 9 | CIDCA 8113, CIDCA 8116, CIDCA 8118, CIDCA 81111, CIDCA 8153, CIDCA 8154, CIDCA 81104, CIDCA 81105, CIDCA 9121 | Isolated from kefir grains | Garrote et al.(6); Diosma et al.(7) |
| *Kluyveromyces lactis* | 1 | CBS 2359 | Unable to grow under anaerobic conditions | Snoek & Steensma (12) |
| *Saccharomyces cerevisiae* | 1 | PE-2 | Unable to grow on lactose | Della-Bianca & Gombert (17); Nijkamp et al.(35) |

\* Culture collection strains of NCYC (Norwich, United Kingdom), CBS (Utrecht, The Netherlands), UFV-3 (Universidade Federal de Viçosa, Brazil), CIDCA (Centro de Investigación y Desarrollo en Criotecnología de Alimentos, Argentina) and PE-2 (Prof. Luiz Carlos Basso, University of São Paulo, Brazil).

### 6.2. Serial dilution spotting assay

The dilution spotting assays were based on the protocol of Della-Bianca & Gombert (2013) (17) with modifications. Cells from a single colony were aseptically transferred from cultures on YGC (yeast extract 5 g/L, glucose 20 g/L, chloramphenicol 0.1 g/L, agar 14.9 g/L)-plates to a 5 mL pre-culture in YPD medium and grown for 24 h at 30 °C, 200 rpm. Cells from this cultivation were used to inoculate tubes containing 5 mL fresh YPD medium, with an initial absorbance at 600 nm (Abs_600_) of 0.1. Strains were incubated at 30 °C, 200 rpm and grown for 3 h (until early exponential phase). Cells were harvested, centrifugated at 5000 g x 5 min and the pellet obtained was resuspended in sterile water to Abs_600_ 0.1 and four successive dilutions (10^−1^, 10^−2^, 10^−3^ and 10^−4^) were prepared. Five μL of each dilution were spotted with a pipette onto plates, which were incubated for at least 72 h. Plates were prepared with solidified industrial substrates (whey or whey permeate) containing lactose 10% (w/v) and incubated at 37 °C. Different stress conditions were tested: anaerobic incubation, ethanol concentration and pH 2.5. For ethanol stress, the medium was supplemented with 5% ethanol (v/v). For low pH stress plates, the medium was adjusted to pH 2.5 with sterile H_2_SO_4_ 1 mol/L. Anaerobiosis was achieved by incubation of plates in an anaerobic jar with Anaero-Pack CO_2_ generator (Mitsubishi Gas Chemical Company, Japan).

### 6.3. Miniaturized industrial fermentation system

This experiment was performed as described by Raghavendran *et al*. (2017) (13). Cells from one colony of the desired yeast strain were aseptically inoculated in 500 mL baffled flasks containing 100 mL of YPD medium (with 4% glucose) and incubated at 30 °C and 200 rpm. Two industrial media were tested in this system: cheese whey and whey permeate, obtained from the dairy industry in powder form and resuspended to 10 % (w/v) in distilled water. After 12 h, the growing culture was transferred to a 2-L flask containing 1 L of autoclaved diluted cheese whey or permeate (10% lactose w/v) for biomass propagation under static conditions at 30 °C. After total sugar consumption (about 36 h), cells were centrifuged (5000 g x 5 min) and approximately 3.6 g of wet biomass was added to a conical tube (50 mL), previously weighed. The wet biomass was suspended in 2 mL of supernatant from a previous cultivation (free of yeast) plus 6 mL of water, representing the “vat foot” (or *pé-de-cuba*) used in the Brazilian 1G ethanol production process. At times zero, two and four hours, 9.25 mL of industrial medium were added to the conical tubes, simulating the fed-batch process or a single addition of 27.75 ml was performed, for a batch process. Every 2 h, the tubes were gently shaken to remove the trapped CO_2_ and weighed. This was performed during 10 h and the experiment was carried out at 30 °C, 34 °C, or 37 °C. After each fermentation cycle, the tubes were left on the bench overnight. In the following day, the tubes were weighed, homogenized and 1 mL of sample was taken for microbiological analysis (viability and contamination). After that, the tubes were centrifuged (5000 g x 5 min) and the supernatant (wine) was saved at −20°C for HPLC analysis. The pelleted yeast biomass was weighed and treated with a 1 mol/L sulfuric acid solution (just enough volume to achieve pH 2.5) during 1 h. Finally, the tubes were centrifuged (5000 g x 5 min) and the acid solution was withdrawn for “vat foot” preparation and start of the next fermentation cycle.

This procedure was repeated for each fermentation cycle. Experiments were performed in duplicate. Calculation of the ethanol yield was performed as described by Raghavendran *et al*. (2017) (13), employing a correction factor for high cell density cultivations. Cell viability was further evaluated with a dye-based method: methylene blue staining and counting in Neubauer chamber.

### 6.4. Resistance to simulated gastric and intestinal conditions

The simulated gastric and intestinal fluids solutions were prepared as described by Grimoud *et al.* (2010) (29). Yeast culture in stationary phase were washed twice with phosphate-buffered saline (PBS; in w/v: 0.014% KH_2_PO_4_, 0.9% NaCl, 0.08% Na_2_HPO_4_, pH 7.2) and resuspended in simulated gastric fluid (NaCl 125 mM, KCl 7 mM, NaHCO_3_ 45 mM, pepsin 3 g/L) at pH 2.5 and a final yeast concentration of 10^7^ CFU/mL. Suspensions were incubated at 37 °C with stirring at 200 rpm for 90 min. After incubation, cells were washed with PBS and the pellets were resuspended in simulated intestinal fluid (pancreatin 0.1% w/v, bovine bile salts 0.15% w/v) at a final pH adjusted to 8.0. Suspensions were incubated at 37 °C with stirring at 200 rpm for 3 h. Yeast viability was assessed by plating samples collected after incubation in simulated gastric fluid and after incubation in simulated intestinal fluid on YGC-agar.

### 6.5. In vitro characterization of the immunomodulatory properties using the Caco-2-ccl20:luc reporter system

Human colonic epithelial cell line Caco-2 stably transfected with a luciferase reporter construction under control of the CCL20 promoter (designated Caco-2CCL20:luc) were previously described (14). Cells were maintained and routinely grown as described (30). All experiments were performed in serum-free medium in 48-well plates.

Wet biomass of *K. marxianus* CIDCA 9121 obtained from the miniaturized industrial fermentation system was resuspended in a sufficient volume of Dulbecco’s Modified Eagle’s Minimal Essential Medium (DEMEM, GIBCO BRL Life Technologies Rockville, USA) to reach an OD_600nm_ = 0.5 (~10^7^ CFU/mL). Cultured Caco-2 CCL20:luc cells were incubated with the yeast suspension for 30 min. Cells were then stimulated with flagellin (from *Salmonella tyhimurium*, FliC) (1 μg/mL) and incubated for 5 h at 37 °C in a 5% CO_2_ – 95% air atmosphere. All experiments included a basal condition without any treatment, whereas FliC was used as a control for 100% induction of pro-inflammatory response. Then, cells were lysed with Lysis Buffer (Promega, Madison, WI, USA). Luciferase activity was measured in a Labsystems Luminoskan TL Plus luminometer (Thermo Scientific, USA) using a Luciferase Assay System (Promega, Madison WI, USA), as previously described (14). Luminescence was normalized to the stimulated control cells and expressed as percentage of normalized average luminescence ± standard error of the mean (NAL ±SEM) of at least three independent experiments.

### 6.6. Animal experiments

Male BALB/c, 6 weeks old mice with weight over 20 g were specific pathogen-free, provided by *Universidad Nacional de La Plata* animal house. The animals kept in polypropylene cages were maintained under standard conditions, fed with standard laboratory mouse chow and housed in a climate-controlled room on a 12-h light–dark cycle. The experimental protocols were approved by the Animal Ethics Committee of Faculty of Exact Sciences, National University of La Plata, Argentina (Approval No 011-01-15). Before conducting experiments, animals were acclimatized to animal facility conditions for 7 days.

### 6.6.a. Resistance to gastrointestinal conditions in vivo

Mice (5 animals per group) were treated with a suspension of 10^7^ CFU/mL of *K. marxianus* CIDCA 9121 or *K. marxianus* NCYC 1429 (previously grown on cheese whey medium for 24 h) in their *ad libitum* drinking water for 7 days. Control animals received the normal drinking water without any additions. Feces were collected on days 0, 3 and 7; weighed; diluted 100-fold; and resuspended in sterile physiological saline (NaCl 0.85 % w/v). Serial dilutions were performed in sterile physiological saline and yeast colony formation counted after seeding on YGC-agar. The results were expressed as CFU/g of feces. The wash-out of the administered yeast was also determined, repeating the same protocol with feces collected five days after yeast administration had been interrupted. To determine the distribution of the yeast within the gastrointestinal tract, 3 mice per group were sacrificed by cervical dislocation on day 8 of the treatment. One centimeter of each section of the gastrointestinal tract (duodenum, ileum, cecum, and colon) was dissected and the contents scraped from the lumen and resuspended in physiological saline for yeast-colony counts after seeding on YGC agar. The results were expressed as CFU/cm of tissue.

Liver and spleen were aseptically collected and placed in a sterile tube with a volume of YPD broth in order to obtain 1 g organ/10 mL. These suspensions were homogenized, enriched in total viable yeast by incubation for 24 h at 37 °C, and used to inoculate YGC-agar plates. Translocation of yeast was defined by growth of microorganisms on plates after 48–72 h of incubation at 37 °C.

### 6.6.b. Induction of experimental colitis using TNBS

Mice were administered normal water or a *K. marxianus* CIDCA 9121 suspension containing 1×10^7^ yeast/mL during four days, with daily changes of the suspension prior to a single intrarrectal administration of 2.5 mg of Trinitrobenzene sulfonic acid (TNBS, Sigma, USA) in 50% ethanol as vehicle with a final volume of 200 μL. Control animals were inoculated with 50% ethanol in distilled water (8, 31). Animals were sorted in four groups according to administration and inoculation, as follows: Water/Vehicle, Yeast/Vehicle, Water/TNBS and Yeast/TNBS. After 48 h of TNBS inoculation, animals were sacrificed by cervical dislocation and samples of colon were taken for histological analysis (H&E staining) and RNA extraction. Damage was estimated in a double blind manner according to the histopathological activity index (HAI) described by Alex *et al*.,(2009) (31).

### 6.7. Statistical analysis

Comparisons among more than two conditions were analyzed using One-way analysis of variance, (ANOVA), followed by Dunnett’s or Tukey’s post-test. Student’s unpaired t-test and Mann Whitney test were used to determine the significance of differences between two groups. Analyses were performed using GraphPad software version 8.02 (San Diego, CA, USA). *P* values less than 0.05 were considered statistically significant.

## 7. Acknowledgements

MDP is fellow of Argentina National Research Council (CONICET); DER, MR and GLG are members of Scientific Career of CONICET. The authors gratefully acknowledge the financial support provided by FAPESP-CONICET scientific cooperation grants, Consejo Nacional de Investigaciones Científicas y Técnicas (CONICET), Universidad Nacional de La Plata (UNLP), Agencia Nacional de Promoción Científica y Tecnológica (ANPCyT), Fundação de Amparo à Pesquisa do Estado de São Paulo (FAPESP, grant number 2016/50444-0) and Alexander von Humboldt Stiftung for partially funding the research via a Return fellowship to Dr. Romanin. JVM-Jr received a scholarship from FAPESP (grant number 2015/26072-3). AKG and JVM-Jr would also like to acknowledge Dr. Guilherme M. Tavares and Dr. Miriam D. Hubinger for helpful discussions.

